# Variations in kinase and effector signaling logic in a bacterial two component signaling network

**DOI:** 10.1101/2024.11.04.621962

**Authors:** Danielle Swingle, Leah Epstein, Ramisha Aymon, Eta A. Isiorho, Rinat R. Abzalimov, Denize C. Favaro, Kevin H. Gardner

## Abstract

The general stress response (GSR) protects bacteria from a wide range of stressors. In *Alphaproteobacteria*, GSR activation is coordinated by HWE/HisKA2 family histidine kinases (HKs), which can exhibit non-canonical structure and function. For example, while most light-oxygen-voltage sensor-containing HKs are light activated dimers, the *Rubellimicrobium thermophilum* RT-HK has inverted “dark on, light off” signaling logic with a tunable monomer/dimer equilibrium. Here, we further investigate these atypical behaviors of RT-HK and characterize its downstream signaling network. Using hydrogen-deuterium exchange mass spectrometry, we find that RT-HK uses a signal transduction mechanism similar to light-activated systems, despite its inverted logic. Mutagenesis reveals that RT-HK autophosphorylates *in trans*, with changes to the Jα helix linking sensor and kinase domains affecting autophosphorylation levels. Exploring downstream effects of RT-HK, we identified two GSR genetic regions, each encoding a copy of the central regulator PhyR. *In vitro* measurements of phosphotransfer from RT-HK to the two putative PhyRs revealed that RT-HK signals only to one, and does so at an increased intensity in the dark, consistent with its reversed logic. X-ray crystal structures of both PhyRs revealed a substantial shift within the receiver domain of one, suggesting a basis for RT-HK specificity. We probed further down the pathway using nuclear magnetic resonance to determine that the single NepR homolog interacts with both unphosphorylated PhyRs, and this interaction is decoupled from activation in one PhyR. This work expands our understanding of HWE/HisKA2 family signal transduction, revealing marked variations from signaling mechanisms previously identified in other GSR networks.

## Introduction

Bacteria are relatively simple organisms that directly face complex environmental challenges, such as fluctuations in nutrient availability, osmolarity, pH, and temperature. To sense and adapt to their changing surroundings, bacteria commonly use two-component systems (TCSs), minimally comprised of a sensor histidine kinase (HK) and cognate response regulator (RR)^3,4^. Signal input to the HK’s sensor domain modulates autophosphorylation of a conserved histidine residue within its dimerization and histidine phosphotransfer (DHp) domain by an ATP molecule bound to its catalytic ATP-binding (CA) domain. This phosphoryl group is subsequently transferred to a conserved aspartate residue within the receiver (REC) domain of the RR. This relay ultimately affects cellular output, typically via RR-mediated transcription^5–7^.

The vast number of HKs can be classified into families based chiefly on their primary sequence features. Among these groupings, the lesser-studied HWE/HisKA2^8,9^ superfamily is distinguished by a motif near the phosphoacceptor histidine residue (H-box), a conserved arginine (R-box), a long ATP lid, and a glutamate that replaces the first asparagine of the N-box. These unique features evidently result in higher-order differences that further distinguish this family. First, while canonical HKs are membrane-bound and strictly homodimeric, many of the HWE/HisKA2 HKs characterized thus far are soluble and non-dimeric. This phenomenon is exemplified most dramatically by the monomeric EL346^10^ and the hexameric EsxG^11^ proteins. With only two full-length structures solved to date^10,12^, there is still much to be learned about the primary sequence features that underly higher order structural differences and how this family fits into the broader structural picture of sensor HKs.

Another defining characteristic of HWE/HisKA2 family members is their involvement in the general stress response (GSR) networks of *Alphaproteobacteria*^13–20^. The GSR is a gene expression program that enables bacteria to cope with a range of adverse conditions such as oxidative stress, heat shock, and UV exposure^2,13,21^. This response works using a so-called “partner-switching” mechanism, whereby HK activity – and subsequent phosphorylation of a downstream PhyR regulator^22–27^ – controls the activity of a transcriptional inhibitor known as NepR. In the absence of stress, transcription is prevented by NepR binding to the σ^EcfG^ general transcription factor. Stress activates an HWE/HisKA2 HK, phosphorylating the PhyR and promoting sequestration of NepR away from σ^EcfG^, allowing transcription of stress-responsive genes to occur. Details of the mechanism which links PhyR phosphorylation to NepR binding remain unclear, as initial models^25,28^ have proposed that phosphorylation of the PhyR REC domain produces an open state that allows NepR to bind its σ-like (SL) domain, while more recent studies suggest that nascent interaction of NepR with unphosphorylated PhyR precedes PhyR phosphorylation and open state formation^29^.

From this basic architecture, several variations on GSR signaling have been observed, from multiple paralogous copies of various GSR key components^2,13,21,27,30–32^ to differences in the signaling logic of the histidine kinases which control pathway activity. One example of the latter is provided by our prior work on a novel HWE/HisKA2 from *Rubellimicrobium thermophilum* DSM 16684^33^ called RT349^34^ (referred to here as RT-HK). This protein contains a light-oxygen-voltage (LOV)^14,35,36^ sensor domain, which detects blue light via the photoreduction of a bound flavin cofactor and concomitant formation of a covalent protein/flavin adduct. As seen for some of its HWE/HisKA2 relatives, we determined that RT-HK is not strictly dimeric; instead, its dimer-leaning equilibrium shifts towards more compact/monomeric conformations under lit conditions and in the absence of ATP^34^. More unexpectedly, we found that its net kinase activity is higher in the dark than in the light^34^. To our knowledge, RT-HK is the only naturally occurring LOV-HK with its signaling logic inverted from the more standard light-activated mode, though we note that genetic studies of *E. litoralis* DSM 8509 suggest the presence of a dark-activated GSR under partial control of a LOV-HK^37^. A similar inverted logic has been conferred upon some engineered light-sensing HKs via alterations to the helical linkers between the sensor and catalytic domains (stemming from the Jα helix in LOV systems), but no generally applicable pattern for achieving this outcome has been established^38,39^.

Here, we further investigated RT-HK’s oligomeric state and reversed signaling logic, as well as its downstream partners. We found that RT-HK likely uses a signal transduction mechanism similar to light-activated systems, as well as an *in trans* mode of autophosphorylation, and the length and register of its Jα linker can be altered to affect net autophosphorylation levels. Exploring downstream effects of RT-HK, we identified two homologs of PhyR in the genome, RT-PhyR and RT-PhyR′. *In vitro* phosphotransfer measurements showed that RT-HK only specifically signals to RT-PhyR, and at an increased intensity in the dark consistent with autophosphorylation levels. Crystal structures of both PhyR variants uncovered a substantial structural shift in RT-PhyR′ immediately following its phosphorylation site, suggesting a possible mechanism of RT-HK preference for RT-PhyR. Further down the GSR pathway, we observed unexpected interaction modes between the single NepR homolog and both unphosphorylated PhyRs, as the RT-PhyR’:NepR interaction is decoupled from phosphorylation, indicating a phosphorylation-independent function for this homolog. Thus, this system exhibits signaling variations at three levels – HK reversed logic, HK phosphorylation of RT-PhyR′, and binding of RT-PhyR′ to RT-NepR′ – that broaden our view of the signaling paradigms in this class of bacterial two-component pathways.

## Results

### Light signal propagation in a dark-activated histidine kinase

We set out to further investigate the inverted signaling logic and oligomeric state equilibrium of RT-HK we previously identified^34^, seeking to better characterize how RT-HK is able to take the same light input as other studied natural LOV-HKs and return an opposite output. In these systems, the light signal is propagated across a considerable distance of approximately 60 Å from the flavin cofactor within the LOV domain to the ATP in the kinase domain (**Figure 1a**). While the dynamics of light signaling have been examined in some LOV-HKs^40–45^, this is the first time it has been explored in a naturally occurring dark-activated system. We began at the global level, using limited trypsinolysis to probe the accessibility of RT-HK under different illumination conditions (dark vs. lit) and nucleotide states (apo vs. AMP-PNP-bound). Using SDS-PAGE analyses of these experiments, we found that RT-HK is markedly more protease-susceptible under lit than dark conditions (**Figure 1b**). More specifically, quantitation of the intact protein band in these gels (**Figure 1c**) showed that both illumination and nucleotide state influenced protein accessibility to trypsin, with the rank order of protease susceptibility being lit apo > lit AMP-PMP > dark apo > dark AMP-PNP. This trend is reminiscent of that seen for RT-HK oligomeric state, where the equilibrium was shifted towards monomer in the lit apo state and dimer in the dark ATP state^34^.

**Figure 1:**
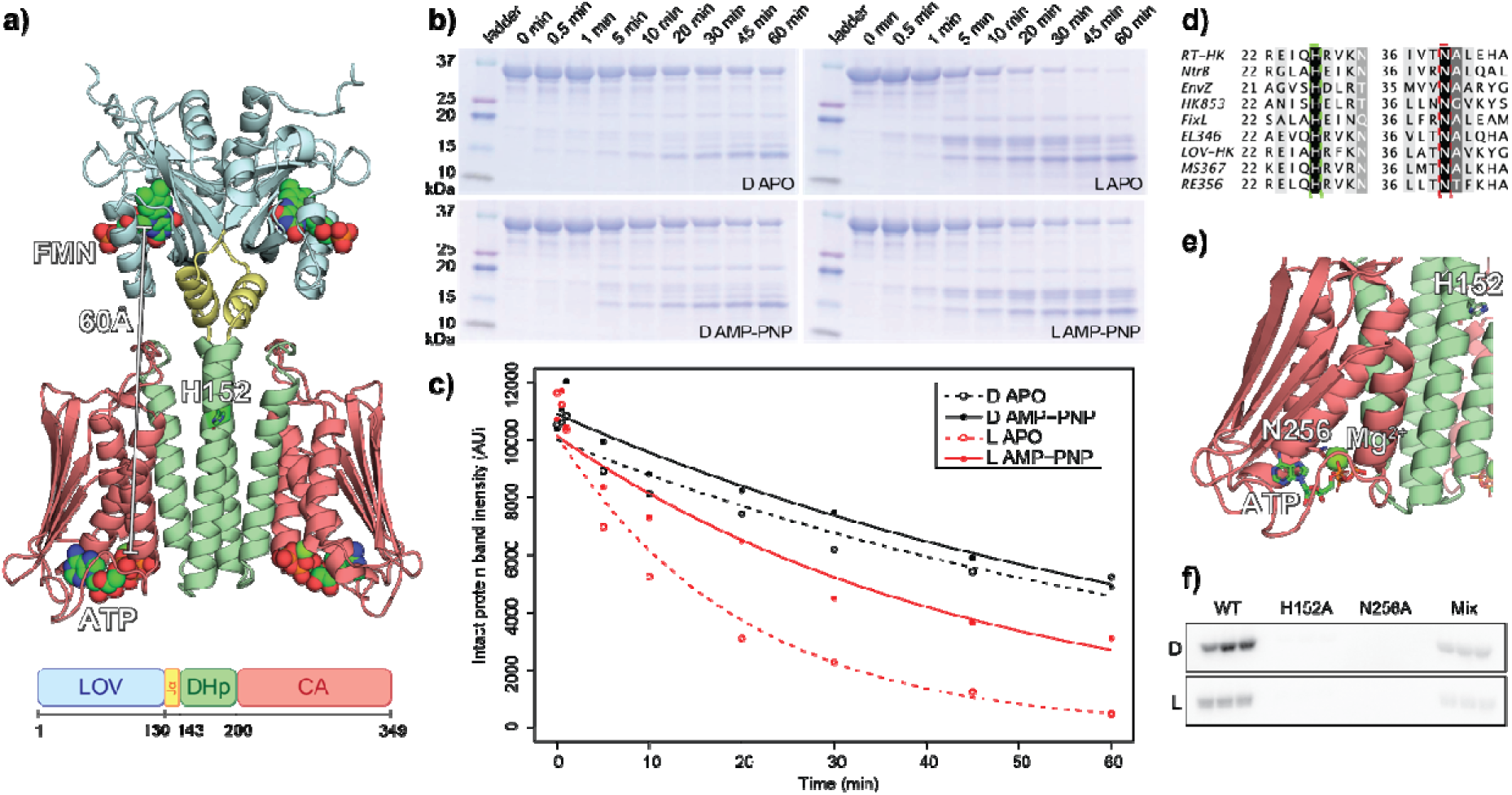
RT-HK protease accessibility is dependent on illumination and nucleotide-binding state, and autophosphorylates *in trans*. **a)** AlphaFold^1^ model of dimeric RT-HK (top) and domain architecture (bottom) and with domains colored correspondingly. The predicted phosphoacceptor histidine residue is shown as sticks, and bound FMN and ATP molecules are shown as spheres, with the distance between them highlighted. **b)** SDS-PAGE gels showing trypsin cleavage pattern of RT-HK over time in dark, lit, apo, and AMP-PNP-bound states. **c)** Intensity of intact protein band over time, with points fit to an exponential curve. **d)** Multiple sequence alignment of RT-HK with other well-studied HKs. Dashed boxes are drawn around the predicted phosphoacceptor His and Mg-chelating Asn residues. **e)** RT-HK AlphaFold model with mutated H152 and N256 residues shown as sticks. **f)** Phosphoimaged SDS-PAGE gels of autophosphorylation assay reactions show substantial ^32^P incorporation only in wild-type (WT) and H152A/N256A mixed (Mix) samples (n=3).

To further elucidate the relationship between RT-HK oligomeric state and activity, we investigated the autophosphorylation mechanism, seeking to establish if this HK phosphorylates its conserved phosphoacceptor His residue using the γ-phosphate of the ATP molecule bound either to the same monomer subunit (*in cis*) or the opposite subunit (*in trans*). The mechanism can be determined by assaying kinase activity on samples composed of a blend of two RT-HK variants which each contain one of two mutants, one deficient in the phosphoacceptor His residue and the other deficient in a conserved Mg^2+^-chelating Asn. Neither of these mutants can autophosphorylate as a homodimer, but when the two are heterodimerized, activity will be restored only if the HK uses a *trans* mechanism^46^. We identified the appropriate RT-HK point mutations needed for this assay, H152A and N256A, by using a multiple sequence alignment^47^ (**Figure 1d**) and predicted structural model^1^ (**Figure 1e**). As expected, each of these mutants alone did not incorporate a measurable amount of ^32^P from the γ-^32^P-ATP substrate in autophosphorylation assays (**Figure 1f**), but we observed restored kinase activity in samples mixing the two mutants. These results indicate that RT-HK autophosphorylates *in trans*. Notably, we saw higher kinase activity of the mixed samples in the dark than in the light, demonstrating that the signaling logic was not affected by the mutations.

The accessibility of RT-HK to hydrogen-deuterium exchange (HDX) was next assessed at the peptide level using a mass spectrometry readout (HDX-MS). Exchange was measured in six different states, varying lit and dark conditions of the LOV domain with either apo, ADP, or AMP-PNP nucleotide states of the kinase domain. Mapping the difference in deuterium uptake between lit and dark conditions in apo, AMP-PNP-, and ADP-bound nucleotide states onto the RT-HK AlphaFold model (**Figure 2**) illustrates several key areas of difference. Intriguingly, peptides throughout the LOV domain exhibited bimodal distributions (**Figure S1**), with an increased ratio of the fast-exchanging population consistently present under lit conditions, regardless of nucleotide state. For two LOV domain peptides with unimodal distributions (17-22 & 70-79) as well as a peptide in the Jα linker helix (127-148), we observed increased exchange under lit conditions. These data are consistent with prior HDX studies of LOV domains and proteins by NMR^48,49^ or MS^50^. The pattern changes substantially in DHp and CA regions, where nucleotide is essential for large-scale HDX. This is exemplified by a peptide from the DHp region (182-189), which experienced similar exchange under dark and lit conditions in the apo state but showed higher exchange in the dark once nucleotide was added. A peptide at the interface between the DHp and CA (213-231) shows nucleotide dependence only in the light. In the ATP lid region of the CA domain (286-315), the peptide shows decreased exchange when AMP-PNP is bound, especially in the dark. Overall exchange in the DHp-CA domains is similar between both nucleotide states, with ADP having a more muted effect. All-in-all, it is clear that nucleotide is required for post-LOV signal transmission and the DHp is at the heart of this process.

**Figure 2:**
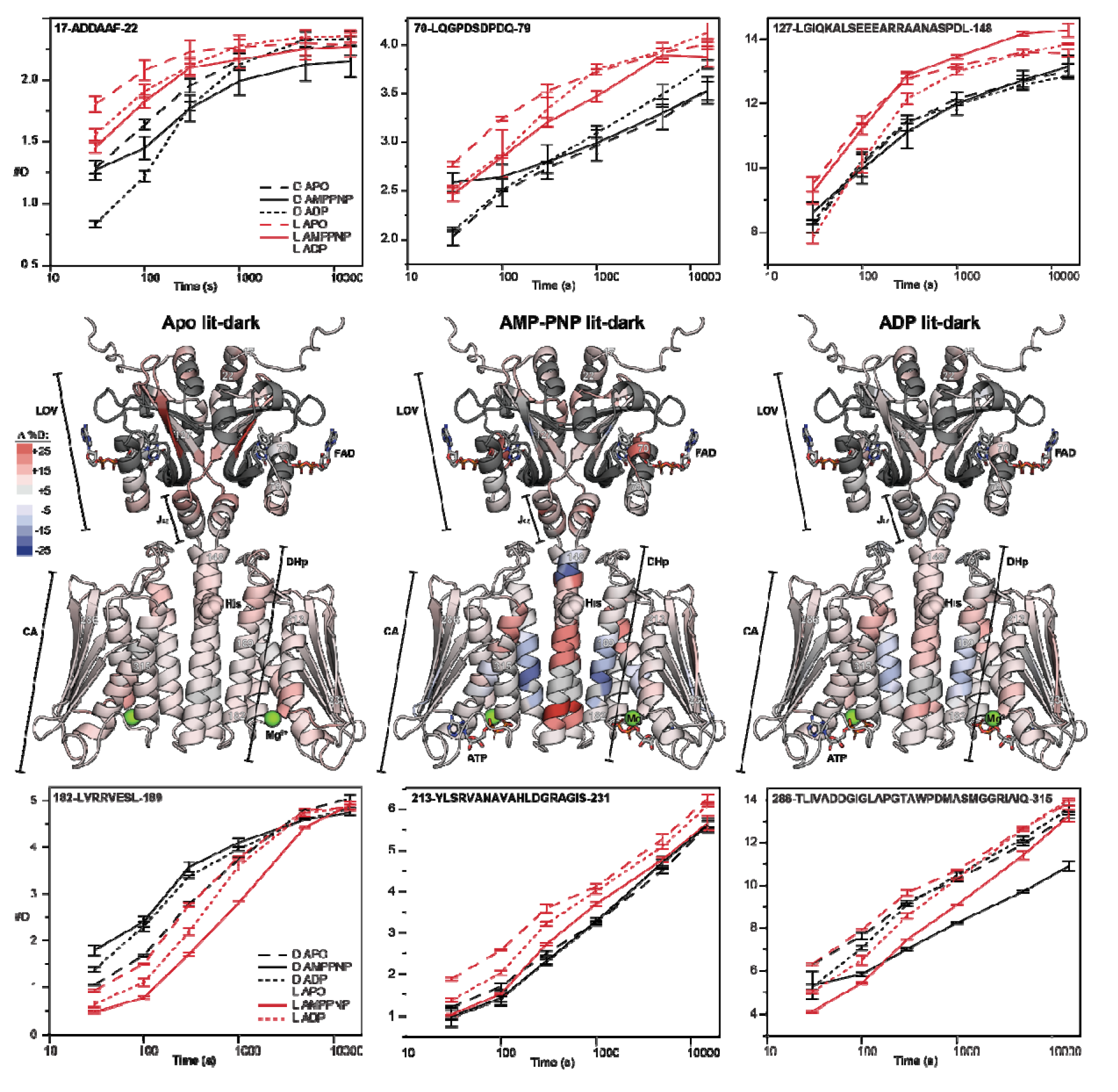
Different regions of RT-HK are differentially affected by light or nucleotide state as assayed by HDX-MS. RT-HK AlphaFold models are colored according to deuterium uptake differences (lit-dark %D) in apo, AMP-PNP-, and ADP-bound states at 300 s, with red (or blue) reflecting increased (or decreased) exchange in the light, respectively. Chiefly bimodal regions are colored dark grey. Uptake plots are shown for selected peptides whose positions are indicated by residue numbers on the models. Error bars represent standard deviation of 3 technical replicates.

Our HDX-MS results highlighted the importance of Jα-containing peptides in signal transduction, prompting us to look further into this region. Previous work on chimeric engineered light-sensing HKs has shown that altering the length of this helix can markedly affect how light input controls net kinase activity^38,39,51^. This phenomenon is generally attributed to conformational changes in the coiled-coil linker between the light-sensing and effector modules of the HK. Intriguingly, the predicted RT-HK structural model has a pronounced break in the coiled-coil between the Jα helix and DHp, likely caused by residue P143 (**Figure 3a**). Four residues surrounding this region were systematically deleted and the effect on net kinase activity was assessed *in vitro* using a γ-^32^P-ATP substrate (**Figure 3b**).

**Figure 3:**
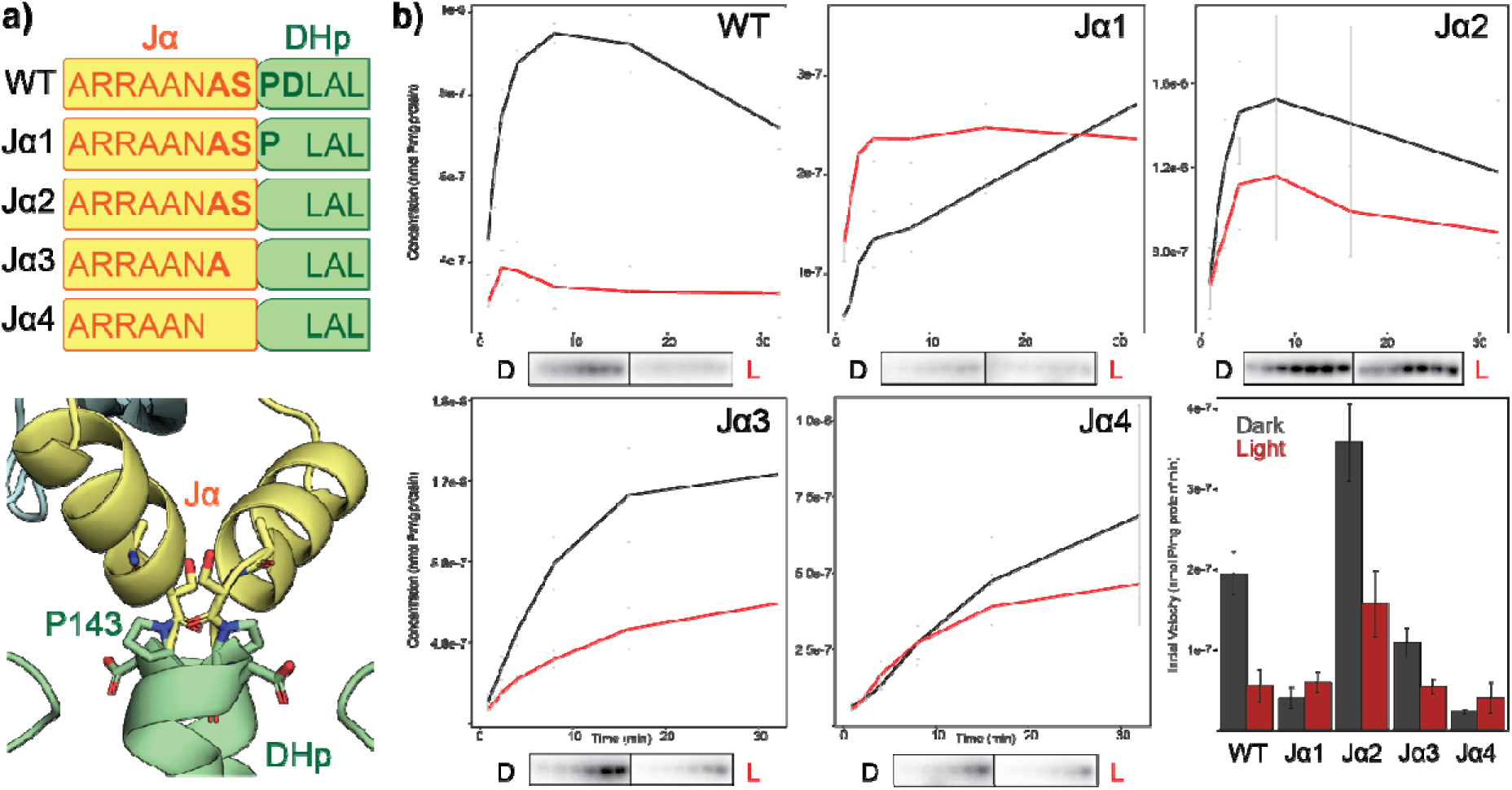
Deletions in Jα linker affect net RT-HK autophosphorylation levels and signaling logic. **a)** Schematic showing positions of amino acid deletions for each construct (top) and AlphaFold model of RT-HK with deleted residues shown as sticks (bottom). **b)** Autophosphorylation assays (plots above, phosphoimages below) are plotted as concentration vs. time for each protein, with dark measurements shown as black lines and lit shown in red. The bar plot shows the initial velocity measured from the linear portion of each curve, with dark measurements shown in dark grey and lit measurements in red. Each point/bar indicates the mean and all error bars span +/-one SD (n=3).

As expected, the wild-type RT-HK showed a sizeable increase in net autophosphorylation in the dark state as compared to the lit state. Surprisingly, removing a single residue to produce the Jα1 mutant caused a drop in dark state activity, resulting in a modest reversal of the signaling logic. The next amino acid deletion (notably, the P143 residue likely “kinking” this region) had the opposite effect – activity in both states increased greatly compared to wild type and the reversed signaling logic was restored. The activity of Jα3 was comparable to wild type, though attenuated in the dark state. And lastly, Jα4 showed a similar effect to Jα1, with a sharp drop in net autophosphorylation levels and another slight reversal of the signaling logic. All-in-all, these results suggest a helical-type periodicity of approximately 3.8 residues, highlighting the importance of the Jα helix and subsequent coiled coil in transmitting signal between sensor and effector domains.

### The role of a dark-activated LOV-HK in a paralogous GSR

After investigating how the light signal propagates through the RT-HK molecule, we set out to identify signaling partner/s and determine if the reversed logic is transferred downstream. Recognizing that HWE/HisKA2 family HKs are often involved in the GSR^13–20^, we searched the *R. thermophilum* DSM 16684 genome for homologs of the key players: the sigma factor σ^EcfG^, the anti-sigma factor NepR, and the anti-anti-sigma factor PhyR response regulator. This search revealed two sets of GSR genes encoded in separate genomic loci, which we refer to as GSR and GSR′ (**Figure 4a**). GSR′ exhibits the typical organization, containing one copy each of NepR, PhyR, and σ^EcfG^ (RT-NepR′, RT-PhyR′, and RT-σ^EcfG^’). Surprisingly, the GSR region also contains its own copies of PhyR and σ^EcfG^ (RT-PhyR and RT-σ^EcfG^), as well as another HWE-family HK (distinct from RT-HK). It is uncommon to find multiple copies of GSR regulators within a single organism and several studies have focused on this phenomenon^27,32,52^.

**Figure 4:**
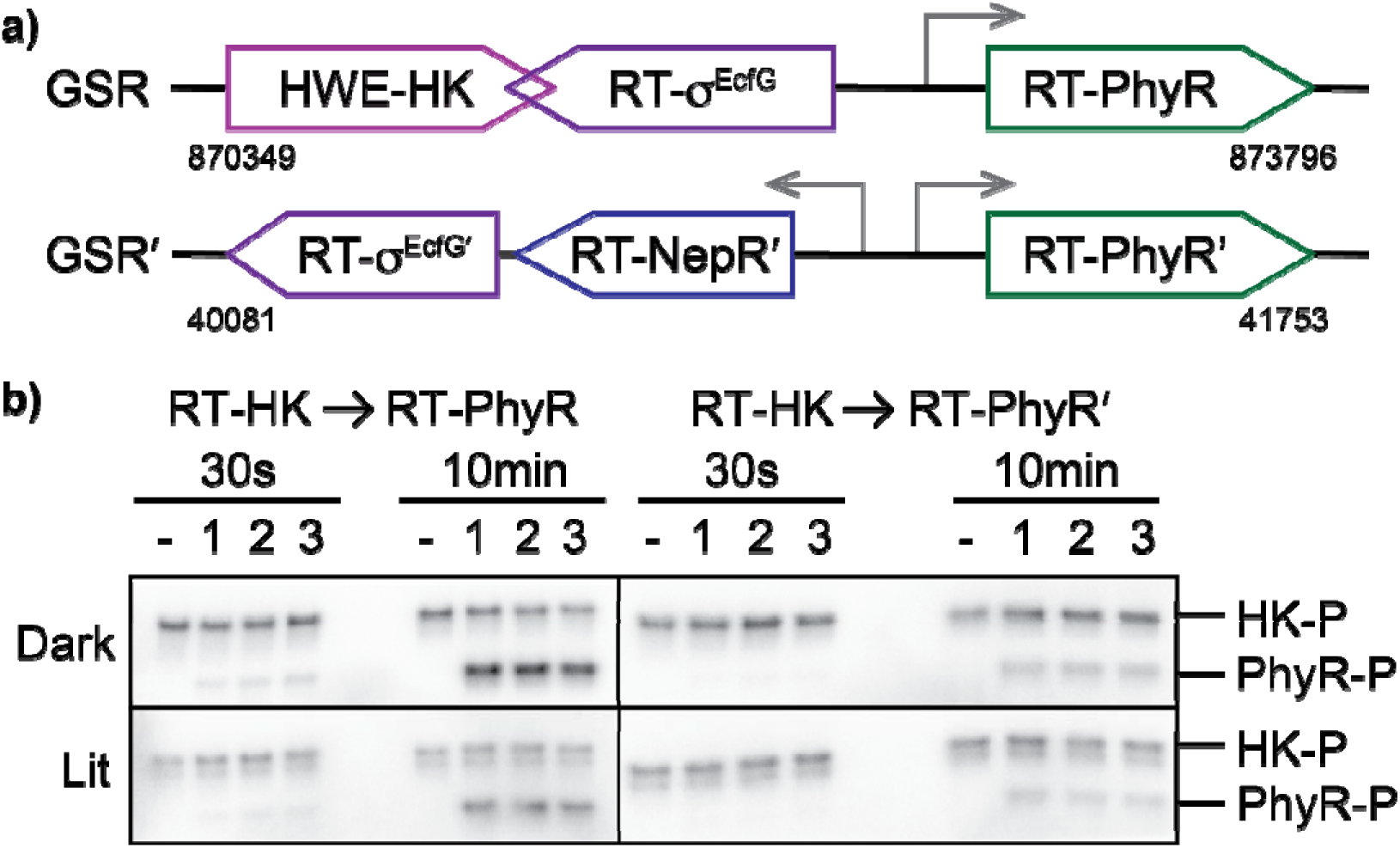
RT-HK reversed signaling logic is transferred to downstream partner RT-PhyR. **a)** Architecture of the two GSR loci in *R. thermophilum* DSM 16684 with genomic locations specified below. Arrows indicate open reading frames. Promoter regions (grey arrows) were predicted from the consensus sequence GGAAC-N_16-17_-C/GGTT^2^. **b)** Phosphoimaged gels tracking the *in vitro* transfer of a ^32^P-labeled phosphoryl group from RT-HK to RT-PhyR (left) and RT-PhyR′ (right). Experiments were performed in dark (top) and lit (bottom) conditions, and samples were taken at 30 s and 10 min timepoints. A negative control (phosphorylated RT-HK only) was run at each timepoint in the left-most lane and triplicate samples were run in consecutive lanes. The top band in each lane corresponds to the mass of phosphorylated RT-HK and bands below this correspond to the mass of the phosphorylated PhyR homolog.

With these two putative downstream PhyR partners in hand, we explored the ability of RT-HK to phosphorylate each using an *in vitro* phosphotransfer assay^53^ (**Figure 4b**). We measured such phosphotransfer at two timepoints: a short timepoint (30 s) to identify the kinetically preferred cognate partner, coupled with a longer timepoint (10 min) to characterize any non-specific transfer. At the short timepoint, we observed transfer only to RT-PhyR, suggesting that it is the cognate partner of RT-HK. Further, the intensity of the phosphorylated RT-PhyR bands were higher in the dark condition at both timepoints, indicating RT-HK’s reversed logic had been transferred downstream. Notably, RT-PhyR′ was phosphorylated only at the longer timepoint and without a marked illumination dependence, suggesting that it is a non-specific partner.

Though there are clear differences between the ability of the two PhyR paralogs to interact with RT-HK, the root of these differences is not immediately apparent from their primary sequences, as they share 64% identity and generally align well with PhyRs from other organisms (**Figure S2**). To provide a structural basis for RT-HK specificity among these two related proteins, we solved the crystal structures of RT-PhyR (1.99 Å resolution; PDB ID: 9BY5) and RT-PhyR′ (2.83 Å resolution; PDB ID: 9CB6); data collection and refinement statistics for both structures are summarized in **Table S1**. RT-PhyR and RT-PhyR′ were solved as a crystallographic trimer and a tetramer, respectively.

In the structures of both RT-PhyR (**Figure 5a**) and RT-PhyR′ (**Figure 5b**), we observed a typical arrangement between the sigma-like (SL) and receiver (REC) domains. The fold of the SL domain is a seven α-helical bundle consisting of the σ2 (α1-3) and σ4 (α5-7) regions, which are connected by a disordered loop that includes α4. Unsurprisingly, no electron density was present for the disordered loop in RT-PhyR′. However, we were able to model this loop into the RT-PhyR structure, where it adopts atypical positions, potentially due to different crystal packing interactions for each chain with neighboring ones (**Figure S3**). The C-terminal end of the SL domain leads into the REC domain, which displays the canonical α/β fold in both proteins.

**Figure 5:**
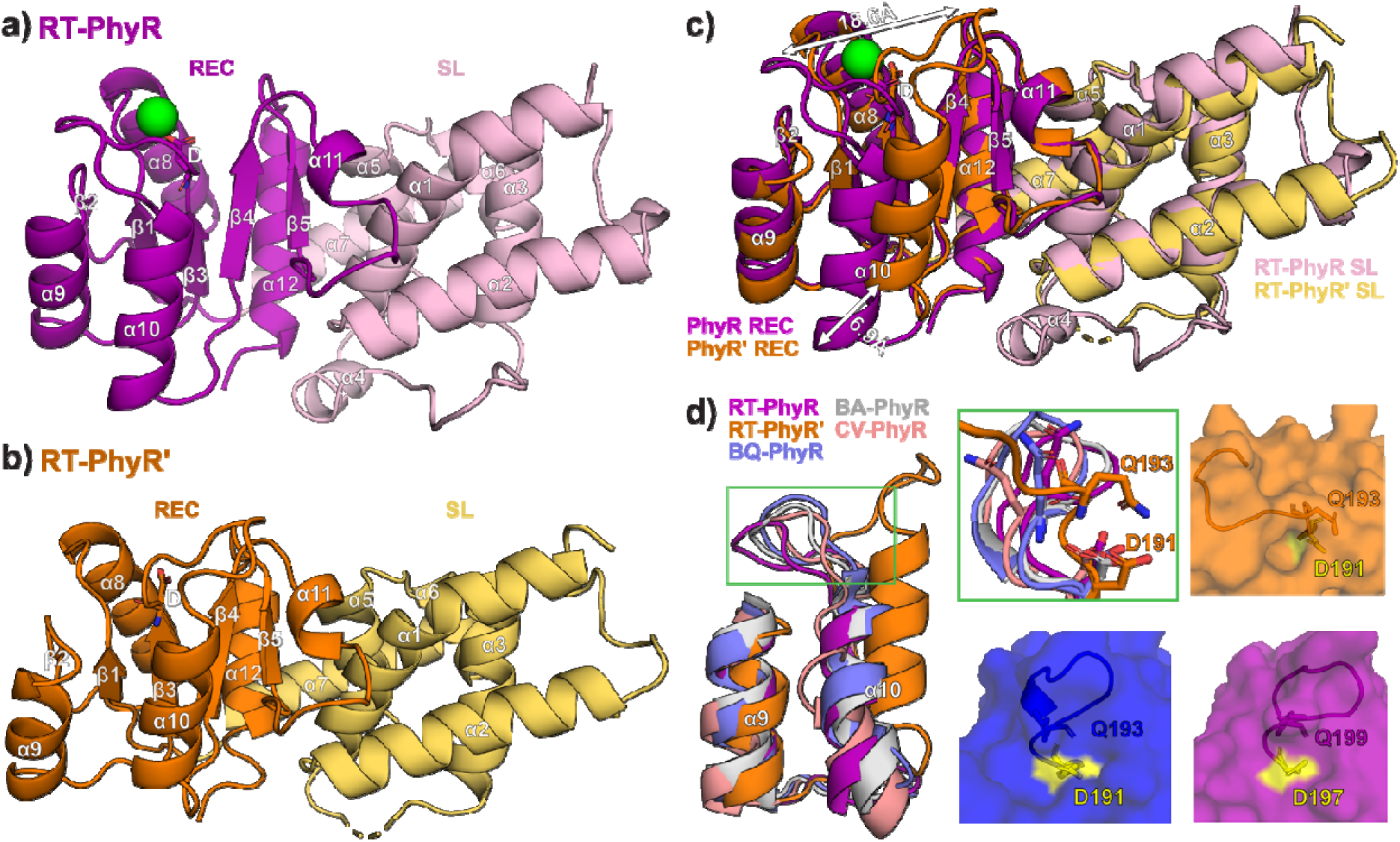
Comparisons of RT-PhyR and RT-PhyR′ crystal structures. **a)** RT-PhyR (PDB ID: 9BY5) and **b)** RT-PhyR′ (PDB ID: 9CB6) crystal structures with domains and secondary structure elements labeled. The phosphoacceptor aspartate residues are shown as sticks. **c)** Aligned RT-PhyR and RT-PhyR′ crystal structures highlight differences in the position of the β3-α10 loop and α10 helix (white arrows). **d)** Aligning several known PhyR structures (labeled according to first two letters of their host organism; BQ PhyR: 5UXW, BA PhyR: 4G97, CV PhyR: 3N0R) to the RT-PhyR/RT-PhyR overlay illustrate that the RT-PhyR′ β3 loop–α10 helix position is the outlier. The green zoomed-in region highlights the unique position of Q193 in RT-PhyR′ near phosphoacceptor D191. Surface representation comparisons illustrate the limited solvent accessibility of D191 in RT-PhyR′.

Aligning the two PhyR paralog structures shows a reasonably high degree of overall similarity (2.7 Å Cα RMSD) (**Figure 5c**), with a notable major shift in the position of the loop connecting the β3 strand (immediately following the phosphoacceptor aspartate) and subsequent α10 helix (α3 in standard REC nomenclature). Adding other PhyR structures into the alignment (**Figure 5d, left**), it is evident that RT-PhyR′ adopts an unusual conformation not routinely seen in other REC domains. This large-scale shift in RT-PhyR′ is likely related to the unique position of a conserved Q residue (Q193), which “leans in” and diminishes the solvent accessibility of the phosphoacceptor D (D191) (**Figure 5d, right**). Two residues in the α9 helix of RT-PhyR′ also adopt distinct positions: a well-conserved R at the N-terminus (R173) and another R at the C-terminus (R184) “reach out” toward the typical position of the β3 loop and α10 helix, respectively (**Figure S4**).

Continuing to the next steps of the signaling pathway, we used ^15^N/^1^H TROSY experiments with uniformly ^15^N-labeled RT-PhyR or RT-PhyR′ to investigate their functional properties and interactions with the single *R. therm.* NepR homolog, RT-NepR′ (**Figure 4a**). When titrated with the phosphoryl group analog BeF_3_^−^, both spectra showed extensive chemical shift perturbations, with slow exchange behavior (**Figure 6a**). RT-PhyR′ reached saturation at the highest BeF_3_^−^ titration point, whereas RT-PhyR did not, suggesting weaker binding by RT-PhyR. Similar slow exchange phenomena were observed when titrating RT-NepR′ to each homolog (**Figure 6b**), but only RT-PhyR′ reached saturation at the highest titration point of RT-NepR′. This suggests that RT-PhyR has a lower affinity for RT-NepR′ than RT-PhyR′ does, similar to the BeF_3_^−^ observations. We underscore that the substantial chemical shift perturbations we observed for RT-NepR′ titrations into the apo-forms (i.e. no BeF_3_^−^) of both PhyR proteins were unexpected, as comparable studies in other PhyR/NepR pairs showed no interaction ^24,25,27^ or much smaller peak shifts^20,29^.

**Figure 6:**
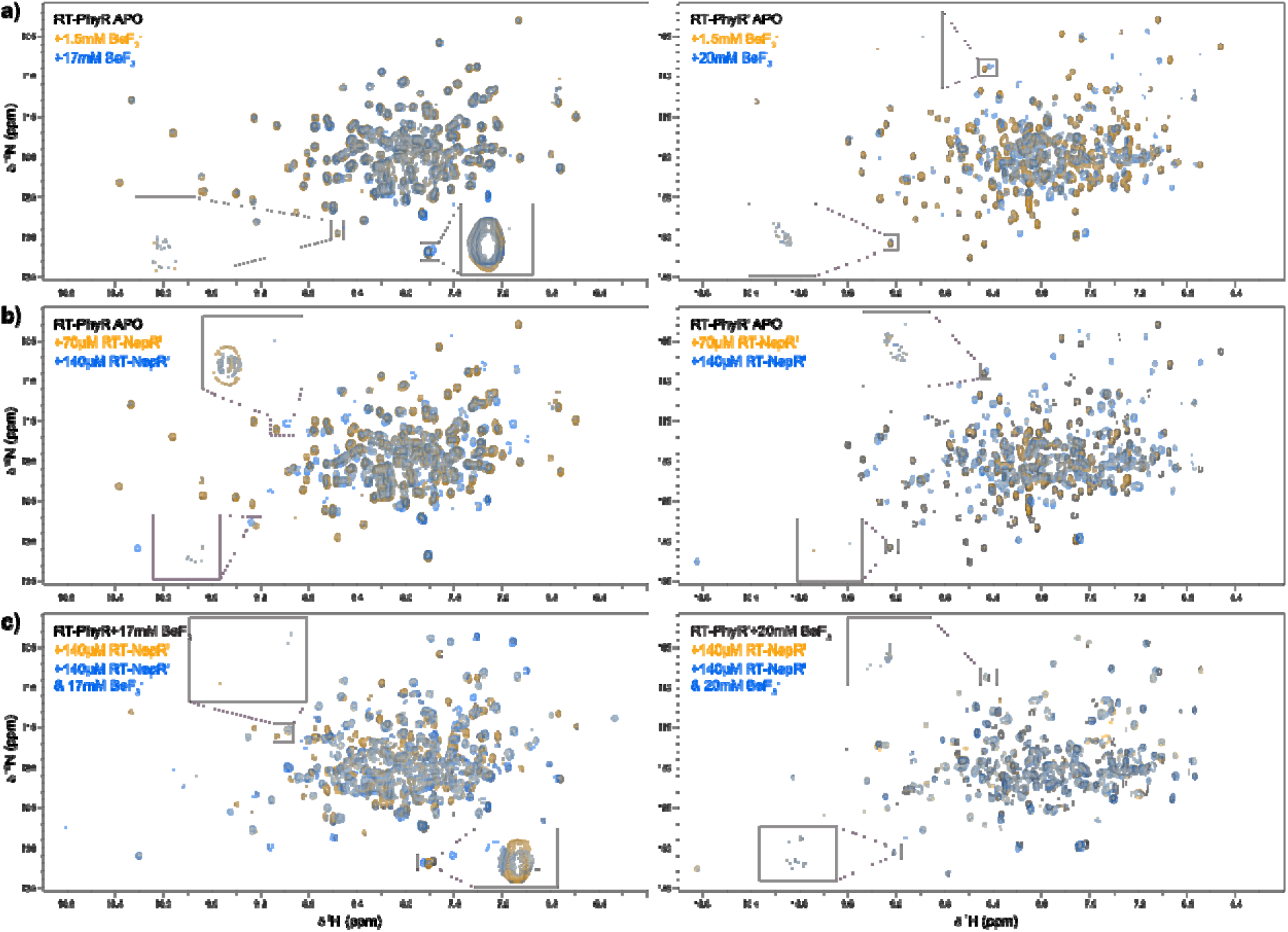
RT-PhyR binds RT-NepR′ and BeF_3_^−^ with positive cooperativity while RT-PhyR′ binds these ligands with negative cooperativity. ^15^N/^1^H TROSY of 210 µM ^15^N-labeled RT-PhyR (left) and 220 µM ^15^N-labeled RT-PhyR′ (right). Proteins were titrated with **a)** BeF_3_^−^, **b)** RT-NepR′, or **c)** both at concentrations indicated in panel insets. Chemical shift perturbations that best illustrate the effects of each condition are expanded in each panel.

To explore the coupling of PhyR proteins binding to BeF_3_^−^ and RT-NepR′, and thus investigate the phosphorylation dependence of this interaction, we titrated BeF_3_^−^ into RT-PhyR or RT-PhyR′ samples that were pre-equilibrated with RT-NepR′ (**Figure 6c**). We observed markedly different results compared to the single titrations above: For RT-PhyR, we saw many peaks shift into positions distinct from those seen when the protein was titrated with either substance alone, indicating a synergistic effect. On the other hand, addition of BeF_3_^−^ to RT-NepR′-equilibrated RT-PhyR′ did not cause any substantial spectral shifts, showing that RT-PhyR′ binds RT-NepR′ to the exclusion of BeF_3_^−^ rather than cooperatively.

## Discussion

In this work, we investigate the unusual properties of RT-HK oligomeric state and signaling logic seen in our prior study^34^ and assess their impact on downstream partners. We uncovered three key features of the *R. therm.* GSR that expand the typical signaling paradigm at different levels: 1) RT-HK is dark-activated, but uses a signal transduction mechanism similar to light-activated systems, 2) RT-HK’s reversed signaling logic is transferred only to RT-PhyR, while RT-PhyR′ is apparently inaccessible to HK phosphodonors, and 3) RT-PhyR′ shows negative cooperativity for activation and RT-NepR′ binding. Our work enhances the current understanding of this complex stress response, introduces novel regulatory modes, and underscores the necessity of testing structural and functional models derived from homology.

To investigate how the inverted signal is propagated through RT-HK, we used HDX-MS (**Figure 2**). Interestingly, bimodal distributions were seen for m/z spectra of peptides throughout the LOV domain, strongly suggesting the existence of two distinct conformational states (**Figure S1**). A higher abundance of the fast-exchanging population under lit conditions suggests that a higher proportion of the LOV domain adopts the associated conformation in the light. At the C-terminus of the LOV domain, in the Jα linker helix, exchange increased under lit conditions in all nucleotide states. This result is consistent with the widely held observation that the Jα helix plays a key role in transmitting light-mediated conformational changes^39,48,51,54^. For example, Jα undocking from the LOV domain upon illumination has been seen in AsLOV2^48,55,56^ and increased accessibility of the Jα helix has been seen in lit-state EL222^49^. Our HDX-MS measurements also highlight that nucleotide is essential for building a light responsive state, as evidenced by changes in the DHp and CA regions upon nucleotide addition. These results are generally comparable to EL346^42^, where light-induced changes are also seen throughout the DHp in the nucleotide-bound states. In both cases, an increase in exchange is seen in the helix surrounding the phosphoacceptor histidine. Overall, RT-HK evidently uses a similar signal transduction mechanism as light-activated systems, despite its reversed signaling logic.

The role of Jα helix properties in LOV-HK signal transmission has been a focus of several studies involving the chimeric engineered YF1 LOV-HK protein^39,51,54^, which also exhibits a “dark active, lit inactive” reversed logic. A common theme among these works is the importance of the heptad periodicity of the continuous coiled-coil linker helix between sensory and output modules in defining signaling logic. Since our structural model of RT-HK shows a proline-mediated break in its analogous linker, we made systematic deletions to alter the length of (and potentially linearize) this region while assessing its role in signal transduction (**Figure 3a**). We saw a clear stepwise change in net dark state activity as single residue deletions were made (**Figure 3b**): Jα1 dropped below WT; Jα2 greatly increased; Jα3 saw a decrease relative to Jα2; and Jα4 exhibited the least activity overall. This pattern is strikingly similar to the periodicity of a coiled coil linker and what has been observed for YF1^39,51,54^.

We next asked how RT-HK’s inverted signaling logic might affect downstream partners. HWE/HisKA2 family HKs are well-established as sensory proteins in the GSR networks of *Alphaproteobacteria*^13–20^. We discovered that the *R. therm.* genome includes two homologous copies of each of the key GSR regulators σ^EcfG^ and PhyR. Many prior studies have focused on the more common occurrence of multiple σ^EcfG^ copies within a genome, but only two have addressed the more rarely-observed systems with multiple copies of PhyR^27,32,57^, and no systemic investigation has been done in a system with the same combination of GSR regulators as *R. therm.*. In general though, the combinations of GSR protein copies and interactions among them tend to vary between systems^31,58,59^; an observation consistent with the hypothesis that multiple copies have diverged to assume different regulatory roles. We used *in vitro* phosphotransfer measurements to assess signaling from RT-HK to both PhyR proteins, identifying RT-PhyR as the cognate partner (**Figure 4b**). Additionally, more efficient phosphotransfer in the dark state indicates that RT-HK’s inverted logic is transferred to RT-PhyR. We further investigated the structural basis of RT-HK preference for RT-PhyR by solving X-ray crystal structures of both PhyR homologs (**Figure 5**). While both structures adopt typical SL and REC domain folds, the β3 loop-α10 helix of RT-PhyR′ is markedly shifted relative to RT-PhyR and PhyRs from various other organisms (**Figure 5d, left**). At the residue level (**Figure 5d, right**), we observed residue Q193 “leaning-in,” decreasing the solvent accessibility of the phosphoacceptor D and limiting its ability to be phosphorylated by an HK partner. We note that this site is accessible to BeF_3_^−^ in our NMR experiments, leaving open the potential for small molecule phosphodonors like acetyl phosphate to control the system.

Our results also lend insights into interactions further downstream between the PhyR homologs and RT-NepR′. NMR titration experiments showed large chemical shift perturbations upon addition of BeF_3_ ^−^ or RT-NepR′ alone to both PhyRs (**Figure 6a,b**). These results are inconsistent with the broadly-accepted hypothesis that REC domain phosphorylation produces an open state PhyR necessary for NepR binding^22–27^, and instead suggest a more complex mechanism, such as the one previously proposed where nascent NepR binding and PhyR phosphorylation act cooperatively to form the inhibitory PhyR∼P/NepR complex^29^. Indeed, our results for RT-PhyR indicate a synergistic effect expected from such a mechanism (**Figure 6c**). However, the analogous interactions seem to be inverted in RT-PhyR′, which exclusively binds RT-NepR′ when provided with both RT-NepR′ and BeF_3_^−^, suggesting a negatively cooperative function for this homolog. The propensity for the unphosphorylated PhyR homologs to bind RT-NepR′ may be related to the density for the typically flexible α4 helix seen in the RT-PhyR structure. This helix plays a role in NepR binding^23,25,28^, acting as a “molecular doorstop” to prevent NepR displacement by the REC domain^21^, and adopts distinct positions based on the presence of NepR. Alignment of the RT-PhyR SL domain with PhyR structures from other organisms reveals a shift in this region (**Figure S3**) where its α4 helix is situated between the expected NepR-bound and unbound positions. However, we are wary of overinterpreting this particular detail, as intermolecular interactions between RT-PhyR molecules in the crystal involve the α4 helix, so this position may be artificially stabilized.

Taken together, our results support the model depicted in **Figure 7**. In the dark, RT-HK increases its net *in trans* autophosphorylation and signals to its cognate partner RT-PhyR much more efficiently than to RT-PhyR′. Still, both proteins adopt distinct conformations upon addition of a phosphoryl group analog. In the absence of phosphorylation, both PhyRs interact extensively with RT-NepR′, each adopting a third conformation. In RT-PhyR, these two pathways from the unphosphorylated conformation evidently act synergistically to promote formation of the final phosphorylated RT-NepR′-bound complex, as previously proposed for *C. crescentus*^29^. In contrast, the absence of a conformation representing the final inhibitory RT-PhyR′∼P/RT-NepR′ complex strongly suggests a phosphorylation-independent function for this homolog, the details of which remain unclear. Ultimately, *in vivo* confirmation of interactions between GSR proteins and their functional importance will be advantageous, but our biochemical and structural data make strong predictions regarding the differential roles of RT-PhyR and RT-PhyR′. All-in-all, these results supplement the existing variety and degree of interactions among copies of GSR regulators. This undoubtedly reflects a wealth of GSR regulatory modes between organisms which have yet to be fully characterized.

**Figure 7:**
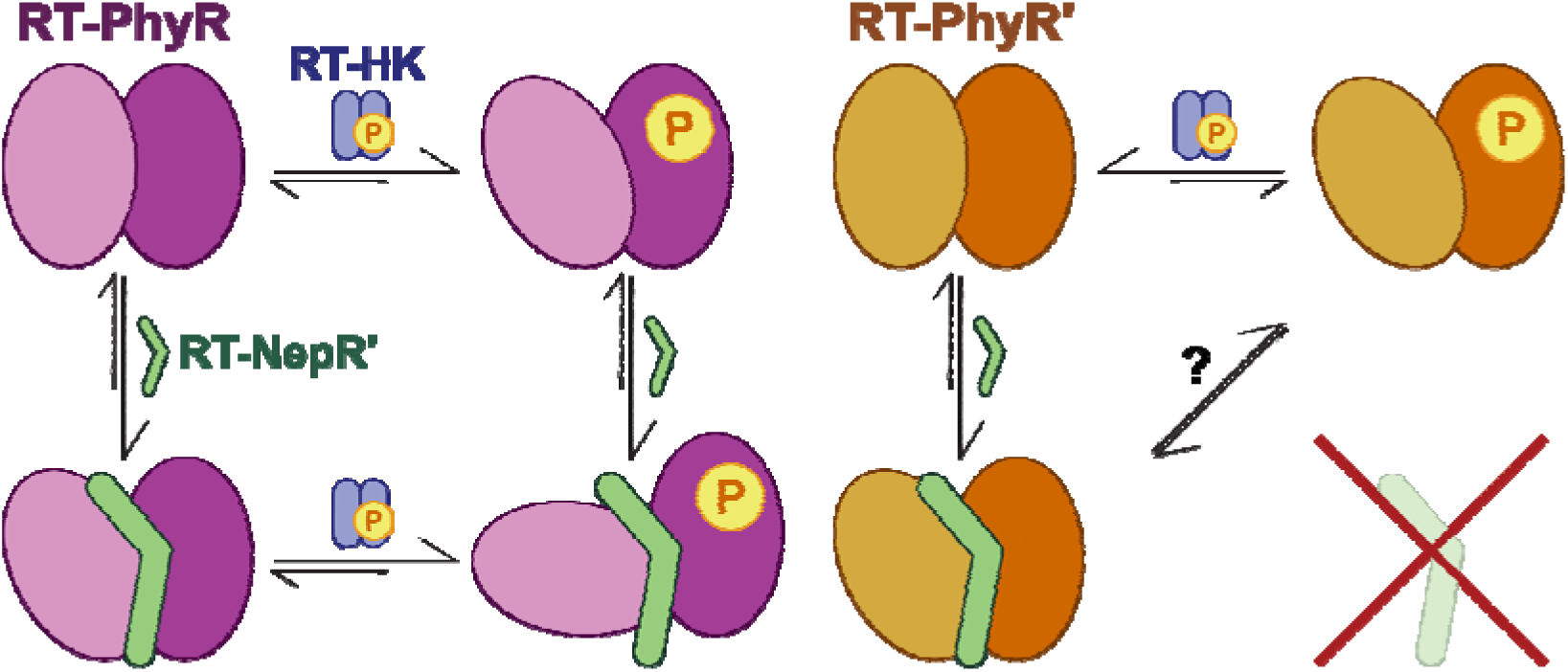
Summary of proposed binding modes for *R. therm.* PhyR homologs. This model illustrates the distinct conformations adopted by RT-PhyR (left) and RT-PhyR′ (right) identified in this study. RT-HK is depicted in blue and RT-NepR′ is in green. The relative sizes of the reaction arrows indicate the equilibrium position.

## Experimental Procedures

### Cloning, protein expression and purification

DNA encoding sequences of RT-HK, RT-PhyR and RT-PhyR′ (NCBI Gene locus tags RUTHE_RS05260, RUTHE_RS05225, and RUTHE_RS12555, respectively) were amplified from *Rubellimicrobium thermophilum* DSM 16684 genomic DNA and Jα deletion genes were ordered from Twist Biosciences. Genes were cloned into the pHis-Gβ1-parallel expression vector^60^. The resulting RT-HK plasmid was used as a template to produce H152A and N256A mutants by site-directed mutagenesis. All constructs were verified by DNA sequencing before being transformed into *Escherichia coli* BL21(DE3) cells (Stratagene). Cells were grown in LB containing 100 µg/mL ampicillin at 37°C and proteins were overexpressed as previously described^42^. Cells were harvested, resuspended in buffer containing 50 mM Tris (pH 8.0), 100 mM NaCl, and 10 mM MgCl_2_, and lysed by sonication on ice. Lysates were centrifuged at 20,000 xg and 4°C for 45 min. Supernatants were filtered through 0.45 µm and bound to a Ni^2+^ Sepharose affinity column (Cytiva) at 4°C. The His6-Gβ1 tagged protein was washed with 4 column volumes of cell resuspension buffer supplemented with 20 mM imidazole and eluted with 250 mM imidazole. Eluted proteins were incubated with His_6_-TEV protease and exchanged into imidazole-deficient buffer by dialysis overnight at 4°C. Proteins were separated from the tags and His_6_-TEV protease by Ni^2+^ affinity chromatography at 4°C and were further purified by size-exclusion chromatography on either a HiLoad 16/600 Superdex 200 pg or a Superdex 200 Increase HiScale 16/40 (Cytiva) at 4°C equilibrated with 50 mM Tris (pH 7.0) (for RT-PhyR, RT-PhyR′, and RT-NepR′) or 10 mM MES (pH 6.5) (for RT-HK and mutants), and 100 mM NaCl, 10 mM MgCl_2_, and 1 mM DTT. For light-sensitive proteins, all purification steps were performed under dim red light. Concentrations were determined from the theoretical absorption coefficient, ε_280_ for PhyRs and RT-NepR’, calculated from the sequence using the ExPASy ProtParam server^61^, and ε_446_ = 11,800 M^−1^ cm^−1^ for all variations of RT-HK.

### Limited trypsinolysis

Reactions were performed on 30 µM RT-HK at room temperature in a buffer containing 50mM Tris (pH 8.0), 100 mM NaCl, 10 mM MgCl_2_, and 1 mM DTT. Samples of apo and 1.6 mM AMP-PNP-equilibrated proteins were equilibrated in lit and dark conditions. Trypsin was added in a 1:1400 ratio to RT-HK, and aliquots were removed to a 4 mM PMSF quench solution at timepoints of 0, 0.5, 1, 5, 10, 20, 30, 45, and 60 min. Samples were subjected to SDS-PAGE analysis and visualized using Coomassie blue stain.

### Production of H152A/N256A heterodimer

Samples of ∼100 µM H152A and N256A mutants in buffer containing 10 mM MES (pH 6.5), 100 mM NaCl, 10 mM MgCl_2_, and 1 mM DTT were mixed and allowed to equilibrate at 4°C for 30 min. The mixture was then added to a dialysis cassette and placed in 500 mL of the same buffer supplemented with 6 M urea and allowed to stir at room temperature for 4 hr. The cassette was then placed into 1 L of the original buffer and stirred overnight at 4°C. The mixture was then reconstituted with 250 µM FMN by 30 min equilibration at 4°C and subsequently separated on a Superdex 200 Increase HiScale 16/40 (Cytiva).

### Autophosphorylation assays of cis/trans and Jα mutants

Experiments were performed as previously described^10,42^. Reactions contained 20 µM protein (40 µM for the heterodimer) in a buffer of 10 mM HEPES (pH 8.0), 100 mM NaCl, 5 mM MgCl_2_, 2 mM DTT, and 10% glycerol. A mixture of unlabeled ATP and 10 µCi [γ-^32^P] ATP was added to each protein to initiate the reaction (final ATP concentration 500 µM). Aliquots were removed at time points of approximately 1, 1.5, 2.5, 4, 8, 6, and 32 min for Jα deletion experiments and 30 min only for cis/trans experiments, then quenched in a 4x SDS-gel loading buffer.

### Hydrogen-deuterium exchange mass spectrometry

30 µM RT-HK was prepared in buffer containing 10 mM MES, pH 6.0, 25 mM NaCl, 5 mM MgCl_2_, and 1 mM DTT and incubated in dark or light conditions with 1.6 mM ADP or AMP-PNP for 30 min. For each time point, one biological and three technical replicates were run. Subsequent labeling and quenching was handled by the automated LEAP HDX platform (Trajan). Labeling was initiated in the same buffer prepared with 100% D_2_O for a precise amount of time before rapid mixing with a chilled quench buffer containing 2 M GdHCl, 3% acetonitrile, and 1% formic acid. Next, an Agilent 1290 pump ran samples through a rapid gradient of 2-90% acetonitrile and 0.15% formic acid at a flow rate of 0.15 mL/min in a Waters Enzymate BEH Pepsin Column. The total digest time was 3 minutes and the resulting peptides were collected and desalted with Hypersil Gold C18 trap 10 mm column and eluted through a Hypersil Gold C18 column (50 mm length × 1 mm diameter, 1.9 μm particle size, Thermo Fisher Scientific) at 0.4 mL/min. Samples were then injected into a Bruker maXis-II ESI-QqTOF high-resolution mass spectrometer. To minimize in-source back exchange, the instrument ion source was lowered to 180°C. Processing of raw mass spectrometry data files was done with Bruker Compass Data Analysis 5.3 and Biotools 3.2 software. Identified peptides were then disambiguated using the PIGEON tool^62^. All data files were then imported into version 3.3 of the HDExaminer software from Trajan Scientific to calculate exchange rate profiles and standard deviation. Sequence coverage was above 98% for each condition.

### Phosphotransfer assays

Phosphotransfer from RT-HK to RT-PhyR and RT-PhyR′ was measured as described previously^24,53^. Reactions took place in a buffer containing 10 mM HEPES (pH 8.0), 100 mM NaCl, 5 mM MgCl_2_, and 2 mM DTT. A mixture of unlabeled ATP and 24 µCi [γ-^32^P] ATP was added to RT-HK to initiate autophosphorylation (final ATP concentration 500 μM, and final HK concentration 10 µM). This reaction was allowed to occur for 10 min before a negative control aliquot was placed into 4x SDS-gel loading quench buffer, and the rest was mixed with an equal amount of RR candidate (final concentration both proteins 5 µM). Aliquots of this mixture were removed at 30 s and 10 min timepoints and placed into quench buffer. For dark measurements, all steps were performed under dim red light. For lit measurements, the samples were illuminated with a blue LED panel just prior to and throughout the course of the experiment. Samples were subjected to SDS-PAGE analysis, gels were dried and exposed to a storage phosphoscreen, and bands were visualized by phosphoimaging with a Typhoon FLA 9500 (Cytiva).

### Crystallization and structure determination of RT-PhyR and RT-PhyR′

Commercially available NeXtal PEGs and ComPAS suite screens were employed to find suitable conditions. Crystals of RT-PhyR and RT-PhyR′ were optimized and grown at room temperature using sitting-drops vapor diffusion method in the presence of 5 and 10 mM MgCl_2_, respectively. The RT-PhyR crystallization buffer consisted of 16% (w/v) PEG 10000 and 0.1 M Tris (pH 8.5). RT-PhyR′ was crystallized from a solution of 0.2 M KSCN, 20% (w/v) PEG2250, and 0.1 M BIS-TRIS propane (pH 6.0). Resulting crystals were cryoprotected with LV CryoOil (MiTeGen), looped, and flash-cooled in liquid N_2_ prior to data collection. Data were collected at the National Synchrotron Light Source II (NSLS-II) light source at Brookhaven National Laboratory using the FMX (17-ID-2) beamline for RT-PhyR and the AMX (17-ID-1) beamline for RT-PhyR′. Data were processed using the autoPROC toolbox^63^, resulting in datasets at 1.99 Å (RT-PhyR) and 2.83 Å (RT-PhyR′) resolution. Balbes^64^ was used to produce search models, and structures were determined by molecular replacement with Phaser^65^. Several cycles of refinement were conducted using Coot^66^ and Phenix^65^. Final data collection, processing, and refinement parameters are provided in **Table S1**.

### Titration experiments with BeF_3_^−^ and RT-NepR′

Starting samples contained 215 µM ^15^N-labeled RT-PhyR and RT-PhyR′ in 50 mM Tris (pH 7.0), 100 mM NaCl, 10 mM MgCl_2_, and 5% D_2_O. These were titrated with BeF_3_^−^ (using a fresh 400 mM stock solution prepared by mixing 400 mM BeCl_2_ and 1.2 M NaF) to concentrations ranging from 1.5-20 mM, or with RT-NepR′ (using a 220 µM stock solution) to concentrations ranging from 70-140 µM (while RT-PhyR/′ concentrations decreased accordingly). We note that high concentrations of BeF_3_^−^ appeared to precipitate RT-PhyR over time, leading us to use slightly different maximum concentrations of this reagent for data shown in Fig 6 (17 mM for RT-PhyR, 20 mM for RT-PhyR’). ^15^N/^1^H TROSY (^15^N/^1^H-WADE-TROSY^67^) spectra were collected at 313.1K on a Bruker Avance III HD spectrometer equipped with a 5mm TCI CryoProbe and operating at a ^1^H frequency of 800.05 MHz. NMRFx Analyst^68^ was used for data processing and analysis.

## Data availability

Structure factors and atomic coordinates have been deposited in the Protein Data Bank with PDB IDs 9BY5 and 9CB6. The complete MS proteomics data set is available on the PRIDE database under the dataset identifier PXD060849.

## Supporting information

This article contains supporting information, including Table S1 and Figures S1-4.

## Acknowledgements

The authors would like to thank Sean Crosson (Michigan State University) and members of the Gardner Laboratory for useful discussions. We would also like to acknowledge Joseph DiCandia for assistance with figure building and manuscript review. This research used the AMX and FMX beamlines of the National Synchrotron Light Source II, a U.S. Department of Energy (DOE) Office of Science User Facility operated for the DOE Office of Science by Brookhaven National Laboratory under Contract No. DE-SC0012704. The Center for BioMolecular Structure (CBMS) is primarily supported by the National Institutes of Health, National Institute of General Medical Sciences (NIGMS) through a Center Core P30 Grant (P30 GM133893), and by the DOE Office of Biological and Environmental Research (KP1607011).

## Funding and additional information

This work was supported by the National Institutes of Health grants R01 GM106239 (to K.H.G.) and T32 GM136499 (supporting L.E.), and National Science Foundation grant 1852496 (supporting R. Aymon), and the Mina Rees Dissertation Fellowship from the CUNY Graduate Center (to D.S.). The content is solely the responsibility of the authors and does not necessarily represent the official views of the National Institutes of Health.

## Abbreviations

CA: catalytic ATP-binding
DHp: dimerization histidine phosphotransfer
HK: histidine kinase
GSR: general stress response
HDX-MS: hydrogen-deuterium exchange mass spectrometry
LOV: Light-Oxygen-Voltage
NMR: nuclear magnetic resonance
REC: receiver domain
RR: response regulator
SL: sigma-like domain
TCS: two-component system
TROSY: transverse relaxation-optimized spectroscopy

