## Supporting Information for "Variations in kinase and effector signaling logic in a bacterial two component signaling network"

Danielle Swingle[1,2], Leah Epstein[1,2], Ramisha Aymon[1,3], Eta A. Isiorho[1], Rinat R. Abzalimov[1], Denize C. Favaro[1], Kevin H. Gardner[1,3,4*]

[1]: Structural Biology Initiative, CUNY Advanced Science Research Center, New York, NY 10031

[2]: Ph.D. Program in Biochemistry, The Graduate Center – City University of New York, New York, NY 10016

[3]: Department of Chemistry and Biochemistry, City College of New York, New York, NY 10031

[4]: Ph.D. Programs in Biochemistry, Biology, and Chemistry, The Graduate Center – City University of New York, New York, NY 10016

Contents:

• Supporting Table S1

• Supporting Figures S1-S4

**Table S1: X-ray crystallography data collection and refinement statistics**

| | RT-PhyR (ruthe_01174) | RT-PhyR′ (ruthe_02744) |
|---|---|---|
| *Data collection* | | |
| Space group | I 2 2 2 | P 1 $2_1$ 1 |
| Unit cell parameters (Å) | *a* = 113.91 | *a* = 60.101 |
| | *b* = 118.642 | *b* = 149.543 |
| | *c* = 121.088 | *c* = 73.564 |
| X-ray source | 17-ID-2 at NSLS-II | 17-ID-1 at NSLS-II |
| Wavelength (Å) | 0.979338 | 0.920105 |
| Resolution range (Å) | 34–1.985 (2.056–1.985) | 33.85–2.825 (2.926–2.825) |
| Unique reflections | 56823 (5587) | 30306 (2907) |
| Multiplicity | 2.0 (2.0) | 2.0 (1.9) |
| Completeness (%) | 99.92 (99.64) | 99.33 (96.61) |
| Mean $I / \sigma(I)$ | 13.74 (0.96) | 2.78 (0.68) |
| $R_{meas}$ | 0.03541 (1) | 0.1792 (1.1) |
| $CC_{1/2}$ | 1 (0.459) | 0.983 (0.38) |
| *Refinement* | | |
| Reflections used in refinement | 56818 (5587) | 30305 (2907) |
| Reflections used for R-free | 2675 (242) | 1393 (149) |
| Number of non-hydrogen atoms | 5975 | 7503 |
| *Macromolecules* | 5752 | 7458 |
| *Ligands* | 27 | 0 |
| *Solvent* | 208 | 45 |
| $R_{work}$ | 0.2085 (0.3423) | 0.2328 (0.3265) |
| $R_{free}$ | 0.2372 (0.3635) | 0.2717 (0.3418) |
| RMS(bonds) | 0.01 | 0.003 |
| RMS(angles) | 1.34 | 0.64 |
| Protein residues | 749 | 957 |
| Average B-factor | 54.23 | 59.09 |
| *Macromolecules* | 54.36 | 59.20 |
| *Ligands* | 53.28 | |
| *Solvent* | 50.72 | 41.04 |
| Ramachandran plots | | |
| *Favored (%)* | 97.70 | 94.77 |
| *Allowed (%)* | 2.03 | 4.06 |
| *Outliers (%)* | 0.27 | 1.17 |
| PDB accession code | 9BY5 | 9CB6 |

Statistics for the highest-resolution shell are shown in parentheses.

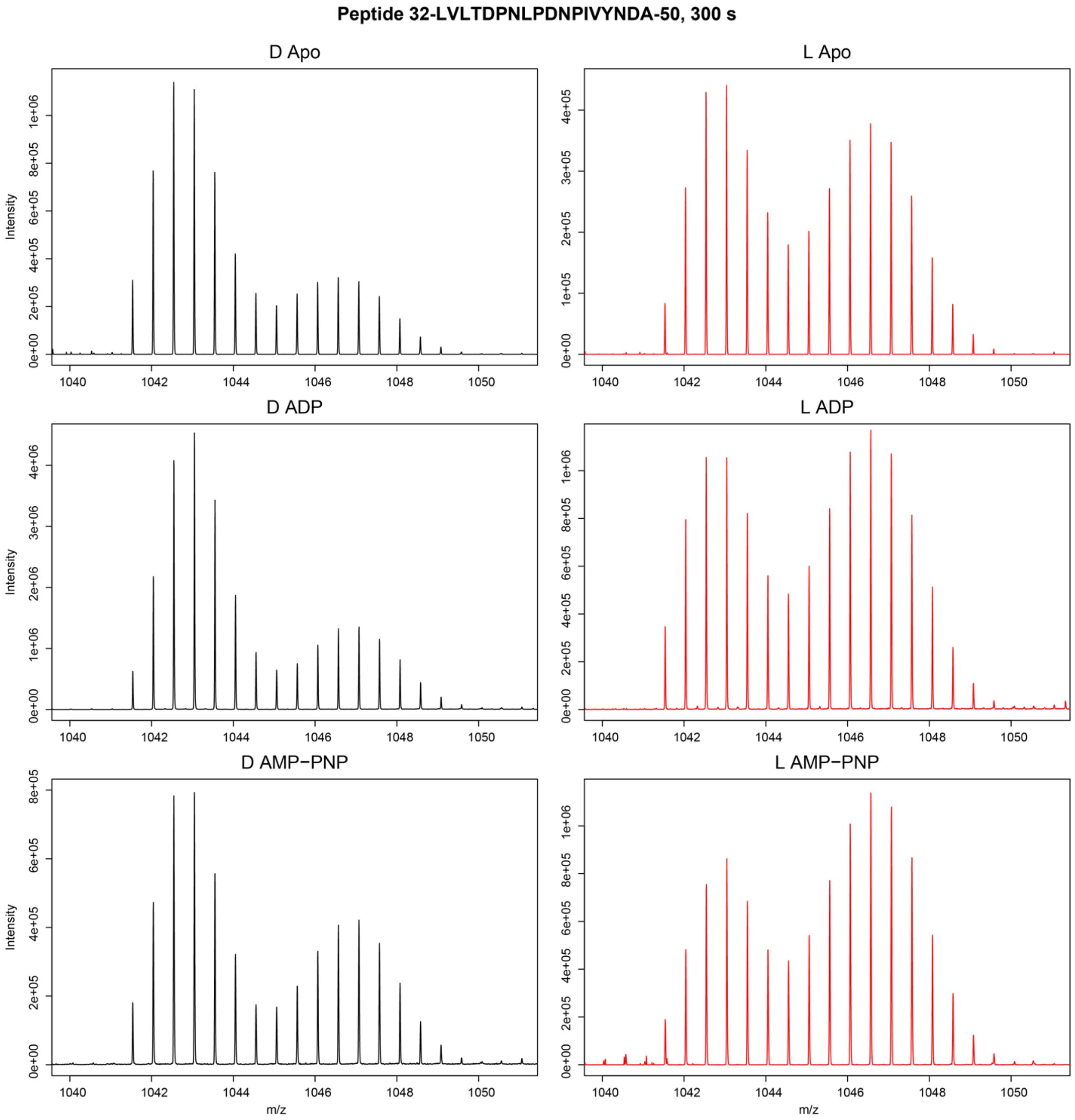


**Figure S1: Lit state peptides have a relatively higher abundance of fast-exchanging population.** A representative example of the bimodal distribution observed for peptides throughout the LOV domain is shown for residues 32-50 in all six protein states. The slow-exchanging population is represented by the left-most peak and fast-exchanging by the right peak. In the vast majority of cases, the fast-exchanging conformation was more abundant under lit conditions.

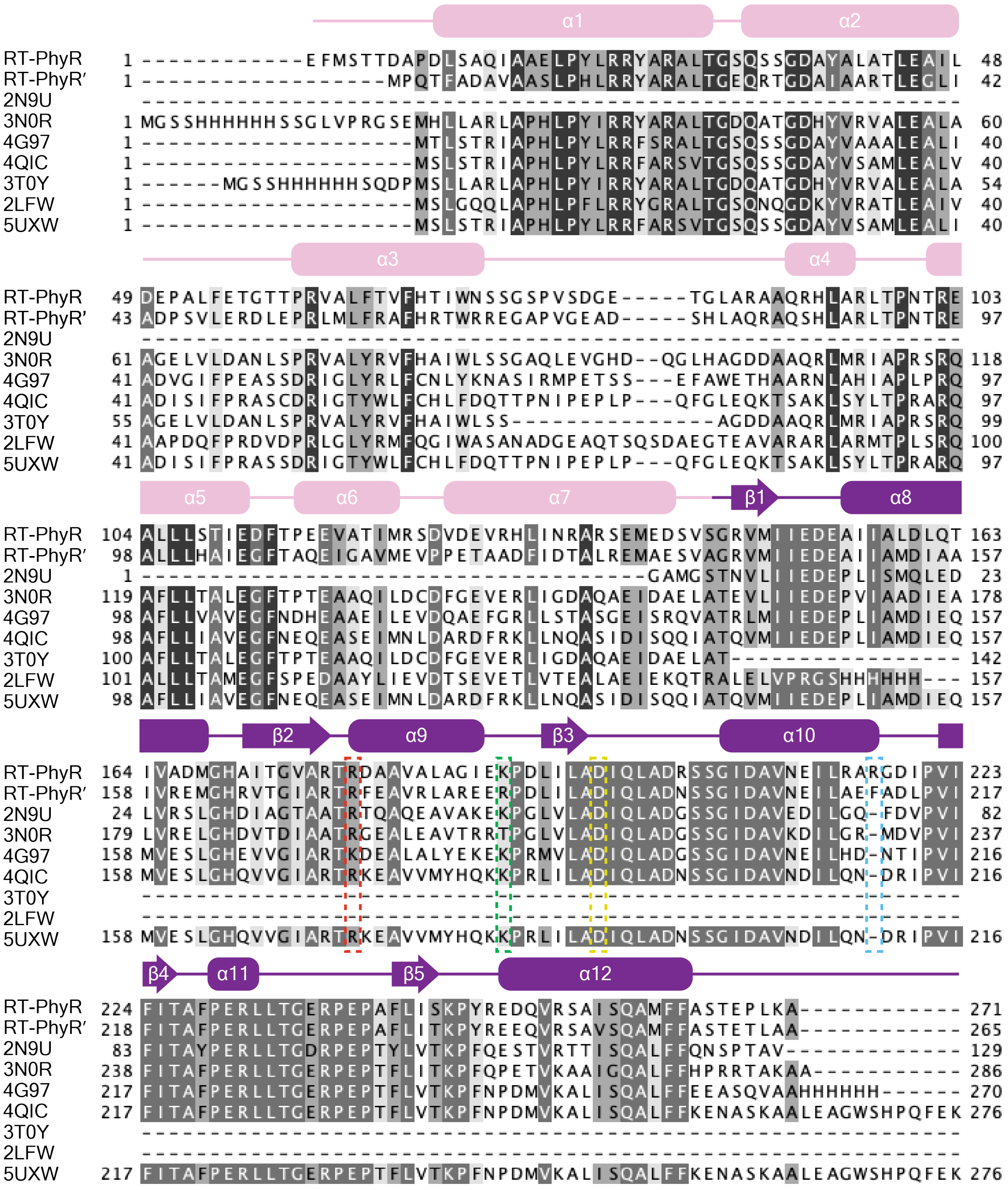


**Figure S2: Multiple sequence alignment of RT-PhyR and RT-PhyR′ with other PhyRs of known structure.** PDB IDs used as identifiers. Residues are shaded by consensus and thresholded at 50%. Secondary structure elements are depicted above each line, with SL domain colored in pink and REC in purple. Dashed boxes indicate important residues: yellow indicates the aspartate phosphorylation site, and red, green, and blue indicate sites defining RT-PhyR′ β3 loop–α10 helix shifts.

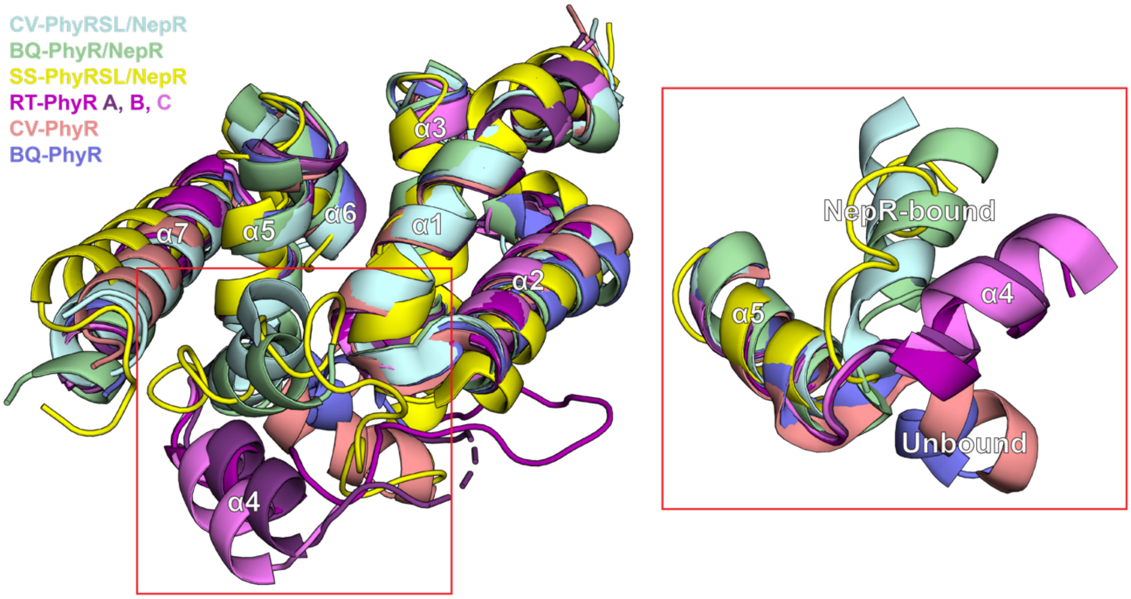


**Figure S3: Position of RT-PhyR α4 differs from all other structures.** Alignment of PhyR SL structures from other organisms with RT-PhyR SL (left). The RT-PhyR α4 helix does not adopt one of the positions defined for NepR-bound or unbound states (right). Structures are labeled according to first two letters of host organism— corresponding PDB IDs are 3T0Y, 4QIC, 2LFW, 9BY5, 3N0R, and 5UXW, from top to bottom. All chains from the RT-PhyR asymmetric unit are shown in different shades of purple, with slightly different α4 locations observed for each chain, which we attribute as likely to different crystal packing interactions with adjacent molecules in the asymmetric unit (chain A) or neighboring ones (chains B and C).

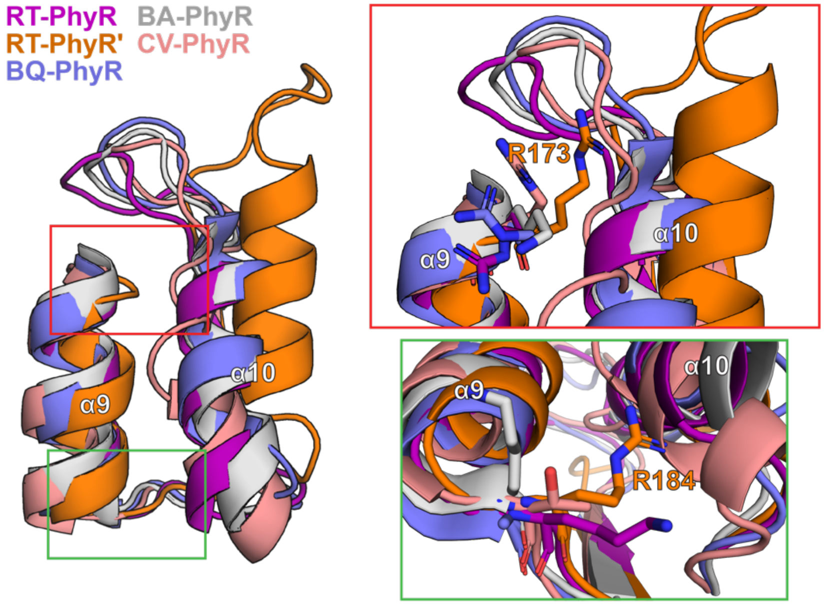


**Figure S4: RT-PhyR′ residue position shifts in α9.** Other known PhyR structures added to the alignment illustrate that the RT-PhyR′ β3 loop–α10 helix position is the outlier. The red zoomed in region highlights the shift in the position of R173 at the top of α9 and the green zoomed-in region shows the shift in R184 at the bottom of α9. Structures are labeled according to first two letters of host organism, with PDB IDs as follows: blue: 5UXW, grey: 4G97, and pink: 3N0R.
